# Single-cell analysis identifies a key role for *Hhip* in murine coronal suture development

**DOI:** 10.1101/2021.02.27.433115

**Authors:** Greg Holmes, Ana S. Gonzalez-Reiche, Madrikha Saturne, Xianxiao Zhou, Ana C. Borges, Bhavana Shewale, Bin Zhang, Harm van Bakel, Ethylin Wang Jabs

**Author notes:** Author contributions: G.H., A.S.G.-R., H.v.B., and E.W.J. designed research; G.H., A.S.G.-R., M.S., X.Z., A.C.B., and B.S. performed research; G.H., A.S.G.-R., X.Z., B.Z., H.v.B., and E.W.J. analyzed data; G.H., A.S.G.-R., H.v.B., and E.W.J. wrote the paper.

## Abstract

Craniofacial development depends on proper formation and maintenance of sutures between adjacent bones of the skull. In sutures, bone growth occurs at the edge of each bone, and suture mesenchyme maintains the separation between them. We performed single-cell RNA-seq analyses of the embryonic, murine coronal suture. Analyzing replicate libraries at E16.5 and E18.5, we identified 14 cell populations. Seven populations at E16.5 and nine at E18.5 comprised the suture mesenchyme, osteogenic cells, and associated populations. Through an integrated analysis with bulk RNA-seq data, we found a distinct coronal suture mesenchyme population compared to other neurocranial sutures, marked by expression of *Hhip*, an inhibitor of hedgehog signaling. We found that at E18.5, *Hhip^-/-^* coronal osteogenic fronts are closely apposed and suture mesenchyme is depleted, demonstrating that *Hhip* is required for coronal suture development. Our transcriptomic approach provides a rich resource for insight into normal and abnormal development.

## Introduction

Craniofacial sutures are critical sites of bone development and growth. During suturogenesis, osteoprogenitors proliferate and differentiate directly from mesenchyme to osteoblasts within the osteogenic fronts (OFs) of adjacent skull bones in a process termed intramembranous ossification, which occurs in the absence of a cartilage template. The OFs are separated by suture mesenchyme (SM), which is maintained for most human sutures into adulthood when they eventually fuse after growth ceases. This process involves multiple interacting signaling pathways, including those for hedgehog (HH), fibroblast growth factor (FGF), ephrin (EPH/EFN), NOTCH, insulin-like growth factor (IGF), retinoic acid (RA), transforming growth factor/bone morphogenetic protein (TGF/BMP), and wingless-related integration site (WNT) signaling (1). Misregulation of such pathways due to genetic or environmental insult can lead to suture dysgenesis, such as wider spacing or premature fusion between bones. Widening of sutures occurs in cleidocranial dysplasia caused by loss-of-function mutations in *RUNX2,* which encodes the master transcription factor for osteogenic differentiation (2, 3). In contrast, loss of suture mesenchyme between bones results in bony fusion or craniosynostosis and reduces the growth potential of that suture. Craniosynostosis is a significant source of human pathology, occurring in approximately 1 in 2,500 births (4). Genetic causes have been identified for approximately 25% of all cases and comprise more than 90 genes involved in a variety of signaling pathways and tissue developmental processes (5–7). The coronal suture, between the frontal and parietal bones on each side of the skull, is fused in approximately 25% of craniosynostosis cases and is the suture most commonly affected in syndromic craniosynostosis cases (6, 8).

The coronal suture is a fascinating suture for study due to its unique biological features. It is the earliest calvarial suture to develop, separating the frontal and parietal bones as they grow from the supraorbital mesenchyme just above the eye (1). In mammals it also lies at the boundary between the neural crest and mesoderm lineages, with the frontal bone derived from neural crest and the parietal bone and SM derived from mesoderm. Little or no mixing occurs between lineages along the length of the embryonic suture (9, 10). The mesoderm extends anteriorly over the neural crest, and as the parietal and frontal bones expand the parietal bone similarly extends to overlap the frontal bone, with a narrow SM separating the bones. In contrast, in the frontal, sagittal, and lambdoid sutures the bones of the calvaria are arranged end-to-end and are separated by a wide SM at embryonic stages.

Various studies have compared RNA expression between human normal and synostotic sutures to identify genes with a role in suture dysgenesis (11). However, these studies often rely on postnatal tissues or bone-derived cells expanded in culture, and may not reflect *in vivo* transcriptomes. Additionally, such studies cannot address how expression of these genes is organized in cell populations. To better understand suturogenesis at the transcriptional and cell population levels with the goal of identifying genes of developmental significance, we have previously applied single-cell and bulk RNA-seq analyses to murine frontal suture at embryonic day (E)16.5 and E18.5 (12). In the current study, we applied these methods to the murine coronal suture. We identified major cell populations comprising SM and OFs of the coronal suture, including populations that differ from those of the frontal suture at the same ages (12).

We found that expression of hedgehog interacting protein (*Hhip*) was enriched in a coronal SM population between E16.5 and E18.5. This was specific to the coronal suture because in the other neurocranial sutures (frontal, sagittal, and lambdoid), *Hhip* expression was enriched in the OFs. *Hhip* encodes an inhibitor of HH ligands and is induced by HH signaling as a component of negative feedback loops regulating the pathway (13). In *Hhip^-/-^* mutant mice, delays in chondrocyte maturation and subsequent ossification of digits, sternum, and vertebrae, which develop from cartilaginous templates in a process termed endochondral ossification, have been described (13), but defects in intramembranous ossification have not been reported. We examined the coronal suture and uncovered a novel phenotype for *Hhip^-/-^* mutants of depletion of the SM. This finding provides new insights into the role of HH signaling in coronal suturogenesis.

## Results

### Single-cell RNA-seq analysis delineates the wild type coronal suture population structure

The coronal suture assumes its definitive morphology of overlapping parietal and frontal bones in the mouse during late embryonic development. This suture remains open during the life of the mouse. We analyzed wild type (WT) coronal suture development by single-cell RNA-seq (scRNA-seq) analysis of four libraries, consisting of two replicates at E16.5 and E18.5. Libraries were derived from strips of coronal sutures spanning the overlapping frontal and parietal bones, including the OFs of each bone and the intervening SM (Fig. 1*A*). Extrasutural tissues were removed to the extent possible to enrich for sutural populations. Cell population identities were determined by reference to published cell type atlases (14–18) and our previously published study of the murine frontal suture (12).

**Fig. 1.**
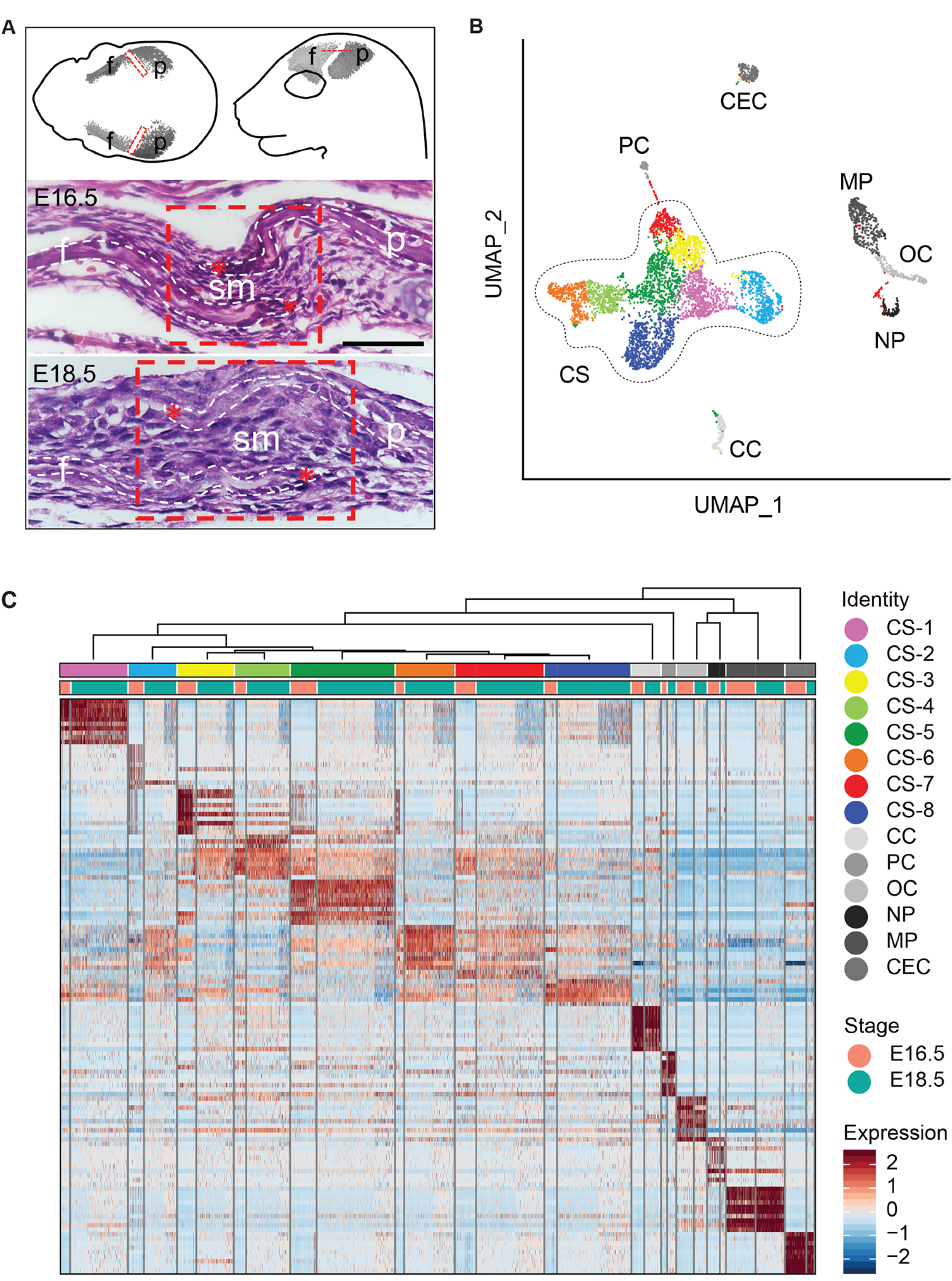
Overview of coronal suture, combined single-cell populations at E16.5 and E18.5. (*A*) Isolation of coronal suture tissue for RNA-seq analysis. At the top left, the schematic shows a top view of the E16.5 mouse head with the mineralized regions of the frontal (f) and parietal (p) bones represented by microcomputed tomography (μCT) images. At the top right, the schematic shows a side view of the E16.5 head. The lower two panels show transverse histological sections of the coronal suture at E16.5 and E18.5, stained with hematoxylin and eosin. Red dashed rectangles, regions between mineralized bone dissected for single-cell analysis; red dashed line, transverse plane of sectioning for histological sections; white dashed outlines, overlapping frontal and parietal bones that are separated by intervening suture mesenchyme (sm); red asterisks, osteogenic fronts. Scale bar, 50 μm. (*B*) Uniform Manifold Approximation and Projection (UMAP) plot of cell-type clusters detected by unsupervised graph clustering of cells from all replicates at E16.5 and E18.5. (*C*) Heatmap of normalized expression of the top ten most significant marker genes for all identified cell populations at E16.5 and E18.5. Columns represent individual cells and rows represent gene signatures enriched (red), depleted (blue), or absent (white) in each cell (FDR ≤ 0.05, lnFC ≥ 0.25).

Analyzing all four libraries combined, we identified 14 cell populations. These included suture mesenchyme and osteoblast populations as well as hematopoietic lineages, vascular cells, and chondrocytes (Fig. 1*B* and *C*). Chondrocytes presumably were derived from the parietal cartilage present at the base of the coronal suture. At E16.5, 56% of cells comprised a supercluster that included suture-specific populations (i.e., suture mesenchyme, osteoblasts) compared to 86% at E18.5.

### Definition and mapping of cell populations at E16.5

At E16.5 the suture-specific supercluster consisted of five populations expressing known markers for two potential mesenchyme populations, osteoblasts, hypodermis, and dura mater (12). These five populations were reclustered separately and seven populations were identified (Fig. 2*A*). To determine the spatial relationships of these seven populations, we used multiplexed single-molecule fluorescence ISH (smFISH) with probes for significant marker genes whose expression levels were high and significantly elevated relative to other populations (Fig. 2*B* and *SI Appendix*, Table S1). Two populations comprised the mesenchyme within the suture. One consisted of SM between the frontal and parietal bones and extending beyond the OFs and was enriched for *Erg, Sox6,* and *Hhip* expression (designated as CS6-4, *<u>c</u>oronal <u>s</u>uture E1<u>6</u>.5 population <u>4</u>*) (Fig. 2*C* and *SI Appendix*, Fig. S1*A*). The other consisted of an ectocranial, or outer, layer of mesenchyme over the frontal and parietal bones and was enriched for *Igfbp3* expression (CS6-6).

**Fig. 2.**
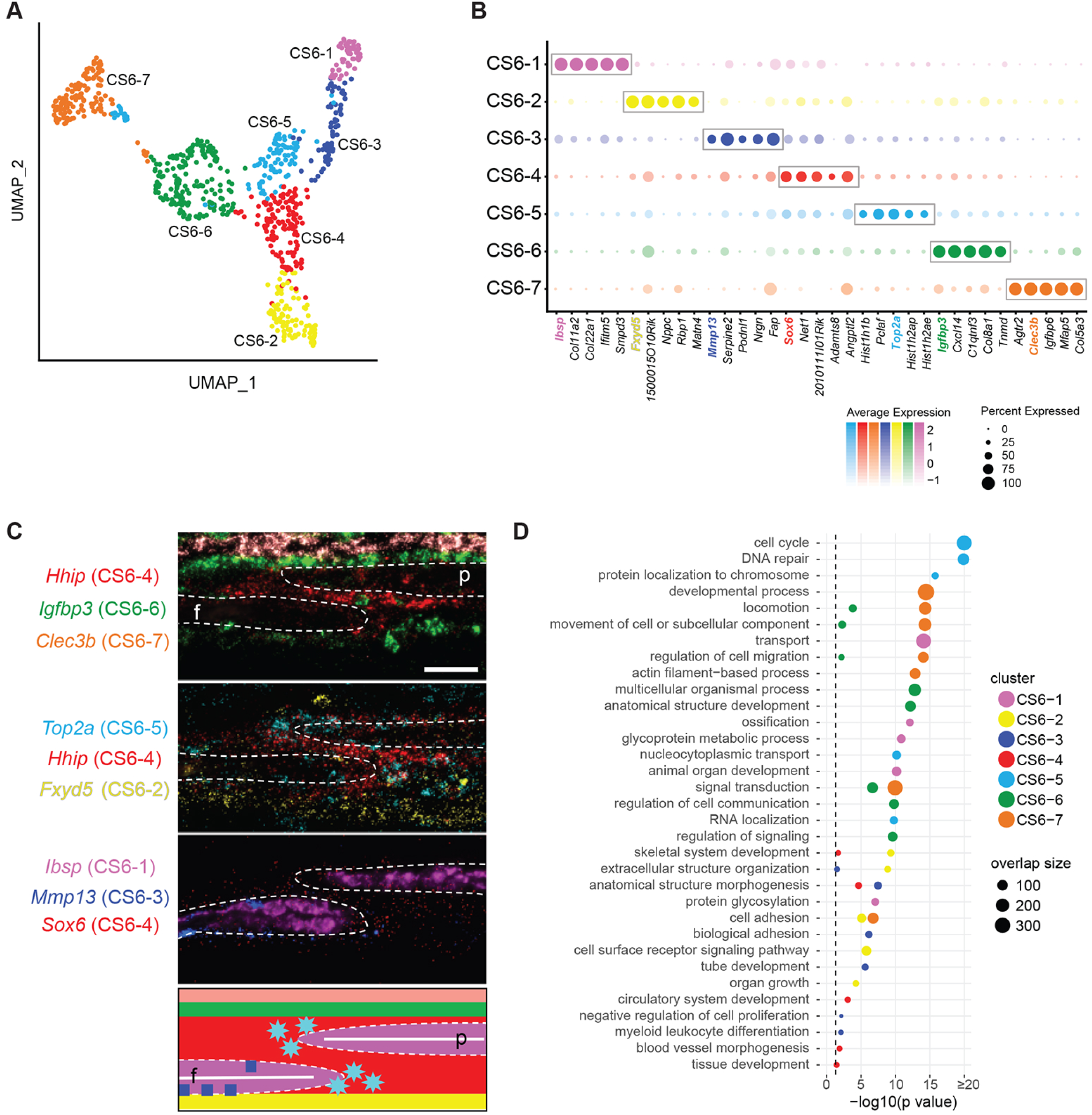
scRNA-seq analysis of the wild type coronal suture at E16.5. (*A*) UMAP plot of cell clusters detected by unsupervised graph clustering of suture-specific populations (i.e., suture mesenchyme, osteoblasts) from two replicates at E16.5. (*B*) Average expression of the top five differentially expressed marker genes (boxed) ranked by FC (FDR % 0.01, lnFC R 0.1) for suture-specific populations. Genes selected for smFISH in (*C*) are colored according to their population membership. (*C*) Localization of populations by smFISH. Each pseudo-colored panel shows an individual section hybridized with probes for the indicated genes. The schematic summarizes the spatial distribution of populations, color-coded as in smFISH. Dashed outlines indicate frontal and parietal bones; white horizontal lines indicate osteoid. f, frontal bone; p, parietal bone. Scale bar, 50 μm. (*D*) Significant GO BP categories of population-specific expression signatures.

Three populations comprised the OFs and osteoblasts. The first consisted of proliferating cells at the OFs of the frontal and parietal bones and was enriched for *Top2a* expression (CS6-5; Fig. 2*C* and *SI Appendix*, Fig. S1*A*). The second consisted of the major population of the frontal and parietal bones and was enriched for *Ibsp* expression, an early osteoblast marker (CS6-1). The third consisted of a smaller population of osteoblasts principally found on the endocranial or inner surface of the frontal bone, enriched for *Mmp13* expression (CS6-3). This was not a frontal bone-specific population, as *Mmp13* also was expressed away from the suture in more mature regions of frontal bones, where high *Mmp13* and *Ibsp* expression were inversely correlated on trabecular bone (*SI Appendix*, Fig. S1*B*). Also in the parietal bone away from the suture, *Mmp13* was expressed on the endocranial surface, while *Ibsp* was expressed on the ectocranial surface (*SI Appendix*, Fig. S1*B*). Two other populations were external to the suture. The hypodermis was above the ectocranial mesenchyme layer and was enriched for *Clec3b* expression (CS6-7). The dura mater was below the suture and was enriched for *Fxyd5* expression (CS6-2).

To characterize the transcriptional programs particular to individual populations we performed Gene Ontology (GO) analysis of the marker genes distinguishing each population identified through differential expression analysis (Fig. 2*D* and *SI Appendix*, Table S1). The main SM population (CS6-4) was enriched for anatomical structure morphogenesis and skeletal system and tissue development. The ectocranial mesenchyme (CS6-6) was enriched for regulation of cell communication, signal transduction, and positive regulation of insulin-like growth factor receptor signaling pathway. The proliferating OF population (CS6-5) was enriched for cell cycle and related processes. The major osteoblast population (CS6-1) was enriched for ossification, collagen fibril organization, protein glycosylation, and protein hydroxylation. The minor osteoblast population expressing *Mmp13* (CS6-3) was enriched for anatomical structure morphogenesis, biological adhesion, and collagen catabolism.

### Definition and mapping of cell populations at E18.5

At E18.5 the suture-specific supercluster consisted of nine populations, which were confirmed by reclustering (Fig. 3*A*). We determined the spatial relationships of these nine populations using smFISH for significant marker genes (Fig. 3*B* and *SI Appendix*, Table S1). Three populations comprised the mesenchyme within the suture. One consisted of the mesenchyme between the overlapping frontal and parietal bones and was enriched for *Hhip* expression (CS8-2; Fig. 3*C* and *SI Appendix*, Fig. S2). The second consisted of mesenchyme beyond the OFs and the region of overlap between the bones and was enriched for *Mest* expression (CS8-4) that also overlapped the *Hhip* expression domain. The third consisted of an ectocranial layer of mesenchyme over the frontal and parietal bones and was enriched for *Col8a1* expression (CS8-7).

**Fig. 3.**
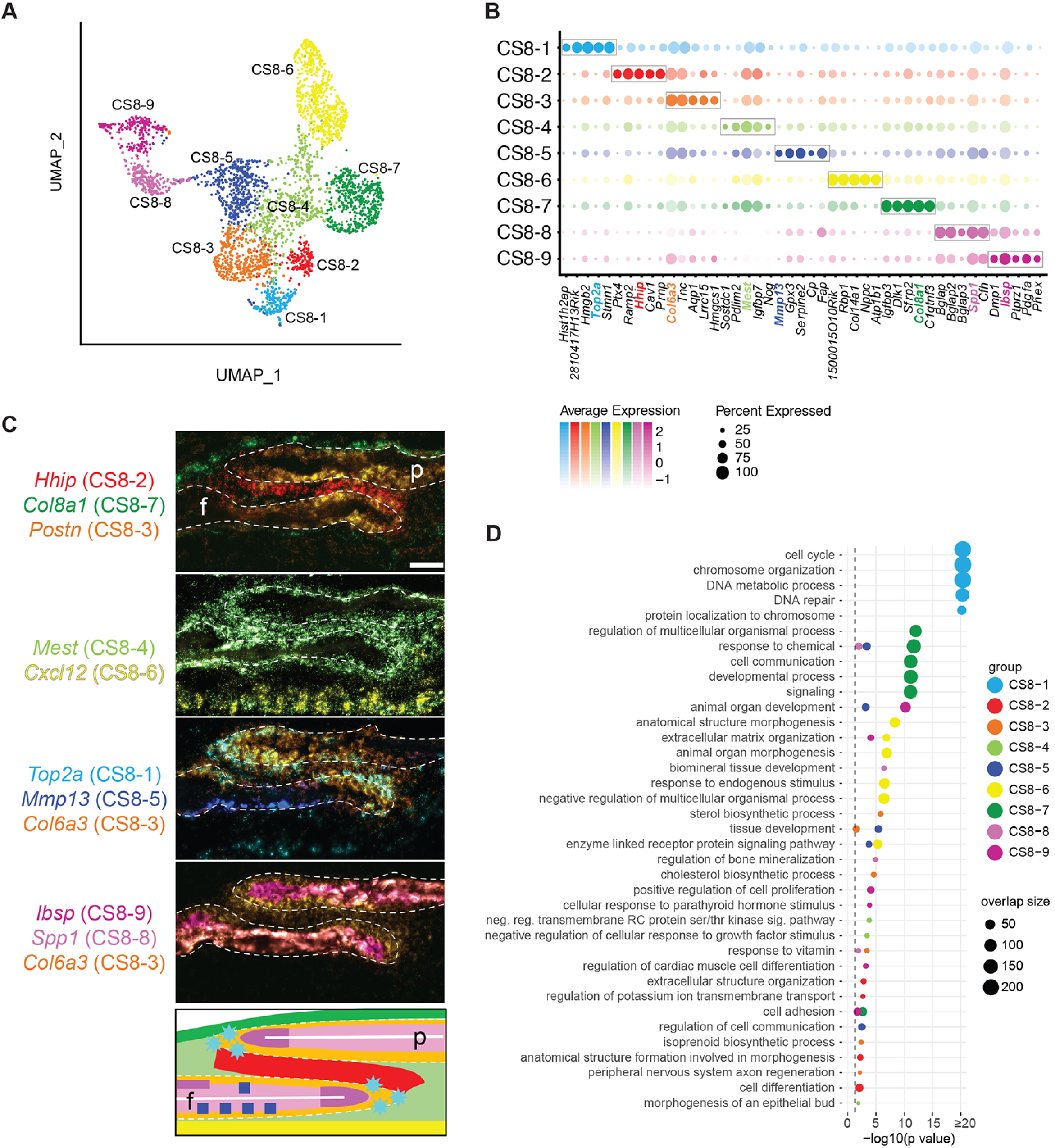
scRNA-seq analysis of the wild type coronal suture at E18.5. (*A*) UMAP plot of cell clusters detected by unsupervised graph clustering of suture-specific populations (i.e., suture mesenchyme, osteoblasts) from two replicates at E18.5. (*B*) Average expression of the top five differentially expressed marker genes (boxed) ranked by FC (FDR % 0.01, lnFC R 0.1) for suture-specific populations. Genes selected for smFISH in (*C*) are colored according to their population membership. (*C*) Localization of populations by smFISH. Each pseudo-colored panel shows an individual section hybridized with probes for the indicated genes. The schematic summarizes the spatial distribution of populations, color-coded as in smFISH. Dashed outlines indicate frontal and parietal bones; white horizontal lines indicate osteoid. f, frontal bone; p, parietal bone. Scale bar, 50 μm. (*D*) Significant GO BP categories of population-specific expression signatures.

Five populations comprised the OFs and differentiated osteoblasts. The first consisted of proliferating cells at the edge of the frontal and parietal bones and was enriched for *Top2a* expression (CS8-1; Fig. 3*C* and *SI Appendix*, Fig. S2). The second was located along the periosteal surfaces, particularly within the region of frontal and parietal overlap and was enriched for *Col6a3* and *Postn* expression (CS8-3). Expression of *Col6a3* appeared stronger than *Postn* towards the OFs. This population also was enriched for expression of *Fgfr2*, which is known to be expressed in OFs and periosteal surfaces of frontal and parietal bones (19–21) (*SI Appendix*, Table S1). The third consisted of osteoblasts extending from the OFs along the osteoid of the frontal and parietal bones and was enriched for *Ibsp* expression (CS8-9). The fourth extended along the bone distal to *Ibsp*-expressing osteoblasts and was enriched for expression of *Spp1*, a marker of more mature osteoblasts (CS8-8). The fifth principally extended along the endocranial surface of the frontal bone and was enriched for *Mmp13* expression (CS8-5). The final population comprised the dura mater below the suture and was enriched for *Cxcl12* expression (CS8-6).

We performed GO analysis of the marker genes distinguishing each population identified through differential expression analysis (Fig. 3*D* and *SI Appendix*, Table S1). For the SM populations, the *Hhip*-expressing SM (CS8-2) was enriched for extracellular structure organization, cell adhesion, cell differentiation, and skeletal system development. The *Mest*-expressing SM (CS8-4) was enriched for negative regulation of transmembrane receptor protein serine/threonine kinase signaling pathway and negative regulation of cellular response to growth factor stimulus. The ectocranial mesenchyme (CS8-7) was enriched for cell communication, cell signaling, and cell migration.

For the OF and osteoblast populations, the GO terms confirmed their diverse roles in skeletal development. The proliferating population (CS8-1) was enriched for cell cycle and related processes. The periosteal population (CS8-3) was enriched for sterol, cholesterol, and isoprenoid biosynthetic processes. The *Ibsp*-expressing osteoblast population (CS8-9) was enriched for extracellular matrix organization and positive regulation of cell proliferation. The *Spp1*-expressing osteoblast population (CS8-8) was enriched for biomineral tissue development. The *Mmp13*-expressing osteoblast population (CS8-5) was enriched for tissue development and collagen metabolic process.

### Population complexity in the coronal suture increases between E16.5 and E18.5

We matched the cell populations at E16.5 with those at E18.5 based on significant expression signature overlaps (*SI Appendix*, Fig. S2*B*). Most SM and osteoblast-related populations were clearly similar between E16.5 and E18.5, including the major *Hhip*-expressing SM populations (CS6-4 and CS8-2), ectocranial SM populations (CS6-6 and CS8-7), *Top2a*-expressing proliferating populations (CS6-5 and CS8-1), *Mmp13*-expressing osteoblasts (CS6-3 and CS8-5), and the dura mater populations (CS6-2 and CS8-6). *Ibsp*-expressing osteoblasts at E16.5 (CS6-1) matched with *Ibsp*-expressing osteoblasts (CS8-9) and a population of more differentiated *Spp1*-expressing osteoblasts (CS8-8) at E18.5. We did not identify discrete populations of *Mest*-expressing SM cells (CS8-4) or *Col6a3*-expressing periosteal cells (CS8-3) at E16.5, but *Mest* and *Col6a3* are expressed at E16.5 in similar domains as at E18.5 (*SI Appendix*, Fig. S1*C*). These differences may reflect changes in the numbers of cells in each of these populations or a lack of sufficient specificity of marker gene expression at this age. *Clec3b*-expressing hypodermis (CS6-7) was not identified at E18.5. This was likely due to easier and more effective removal of extrasutural tissue such as the hypodermis at E18.5 compared to E16.5.

### Ectocranial, periosteal and osteoblast populations are enriched for craniosynostosis genes

Approximately 57 genes have been identified that cause coronal craniosynostosis when mutated in humans (5, 7) (*SI Appendix*, Table S2). We determined the degree of enrichment of these genes among the cell populations at each age. At E16.5 significant enrichments were found in the ectocranial layer (CS6-6; *Cyp26b1*, *Igf1r*, *Lmx1b*, *Stat3*, *Twist1*, *Zic1*; *P* = 0.013) and the *Mmp13*-expressing osteoblasts (CS6-3; *Alpl*, *Atr*, *Fgfr2*, *Fgfr3*, *Kat6a*, *Kmt2d*; *P* = 0.006). At E18.5 significant enrichments were found in the ectocranial layer (CS8-7; *Abcc9*, *Cyp26b1*, *Fbn1*, *Gpc3*, *Igf1r*, *Stat3*, *Twist1*, *Zic1*; *P* = 0.007), the *Mmp13*-expressing osteoblasts (CS8-5; *Alpl*, *Fgfr3*; *P* = 0.044), and the periosteal population (CS8-3; *Fgfr2*, *Flna*, *P4hb*; *P* = 0.029). If we used 96 genes currently associated with craniosynostosis of any suture (5, 7), these five populations also were enriched significantly. In addition, the *Ibsp*-expressing osteoblast population at E18.5 was enriched significantly (CS8-9; *B3gat3*, *Bmp2*, *Fam20c*, *Ihh*, *Irx5*, *Phex*, *Sec24d*, *Sh3pxd2b*; *P* = 0.001).

Approximately 23 genes have been identified that cause coronal craniosynostosis when mutated in mice, of which 11 are models of human disease (22, 23) (*SI Appendix*, Table S2). We determined the degree of enrichment of these 23 genes among the cell populations at each age. At E16.5 significant enrichments were found in the ectocranial layer (CS6-6; *Axin2, Fgf9, Lmx1b, Twist1*; *P* = 0.001) and the *Mmp13*-expressing osteoblasts (CS6-3; *Alpl, Fgfr2, Fgfr3; P* = 0.005). At E18.5 significant enrichments were found in the *Mmp13*-expressing osteoblasts (CS8-5; *Alpl*, *Fgfr3, Runx2*; *P* = 0.0002). If we used 32 genes associated with craniosynostosis of any suture in mice (22, 23), only the *Mmp13*-expressing osteoblast populations were enriched significantly. These two populations were common to the human analysis, but our finding of fewer populations enriched for murine craniosynostosis genes compared to human may be due to the lower number of mouse genes available for analysis. Overall, these enrichment profiles suggest that genes causing craniosynostosis function in different sutural cell populations and thus through varied mechanisms.

### *Hhip* expression distinguishes the coronal suture mesenchyme from other calvarial sutures

Because the molecular processes and cellular composition of suture mesenchyme are poorly characterized compared to those of osteoblast differentiation, we focused on identifying unique features of the coronal suture mesenchyme populations. The coronal suture is morphologically distinct from the other calvarial sutures in that the bones overlap during the embryonic period studied, which may be reflected by a distinct transcriptional signature. We intersected the significant marker genes from all single cell, mesenchymal populations (CS6-4, CS6-6, CS8-2, CS8-4, and CS8-7; *SI Appendix*, Table S1) with our bulk RNA-seq datasets of the calvarial sutures (coronal, frontal, sagittal, and lambdoid sutures; deposited in the FaceBase repository as part of the “Transcriptome Atlases of the Craniofacial Sutures” project; https://www.facebase.org/). These bulk datasets contain SM and OF gene expression profiles of each suture to allow identification of differential gene expression between SM and OFs. This combined analysis of single cell and bulk RNA-seq data identified genes whose average expression was most enriched in the coronal SM compared to enrichment in the OFs (Fig. 4*A*). In this regard, *Hhip*, a marker of CS6-4 and CS8-2, was the most notable as it was predominantly expressed in the SM of the coronal suture whereas it was more highly expressed in the OF of the other calvarial sutures (Fig. 4*A*). We repeated this analysis with an aggregate list of the combined top 10 marker genes of the major SM populations, CS6-4 and CS8-2. *Hhip* was the gene most significantly enriched in coronal SM (Fig. 4*B*). In contrast, other genes such as *Angptl2*, *Adamts8*, *CD34*, *Erg*, *Ets2*, and *Itgb5* were enriched in the SM of all calvarial sutures (Fig. 4*B*). *Hhip* is a component of the HH pathway. To explore this pathway further we made a similar comparison between a list of key members of the HH pathway and bulk SM and OF gene expression in all calvarial sutures. Expression of the Indian hedgehog (*Ihh*) ligand, its receptors *Ptch1* and *Ptch2*, and the HH transcriptional target *Gli1*, was enriched in the osteogenic fronts of all calvarial sutures (Fig. 4*C*). In agreement with this finding, *Ihh* was enriched in single-cell osteoblast populations (CS6-1 and CS8-9; *SI Appendix*, Table S1).

**Fig. 4.**
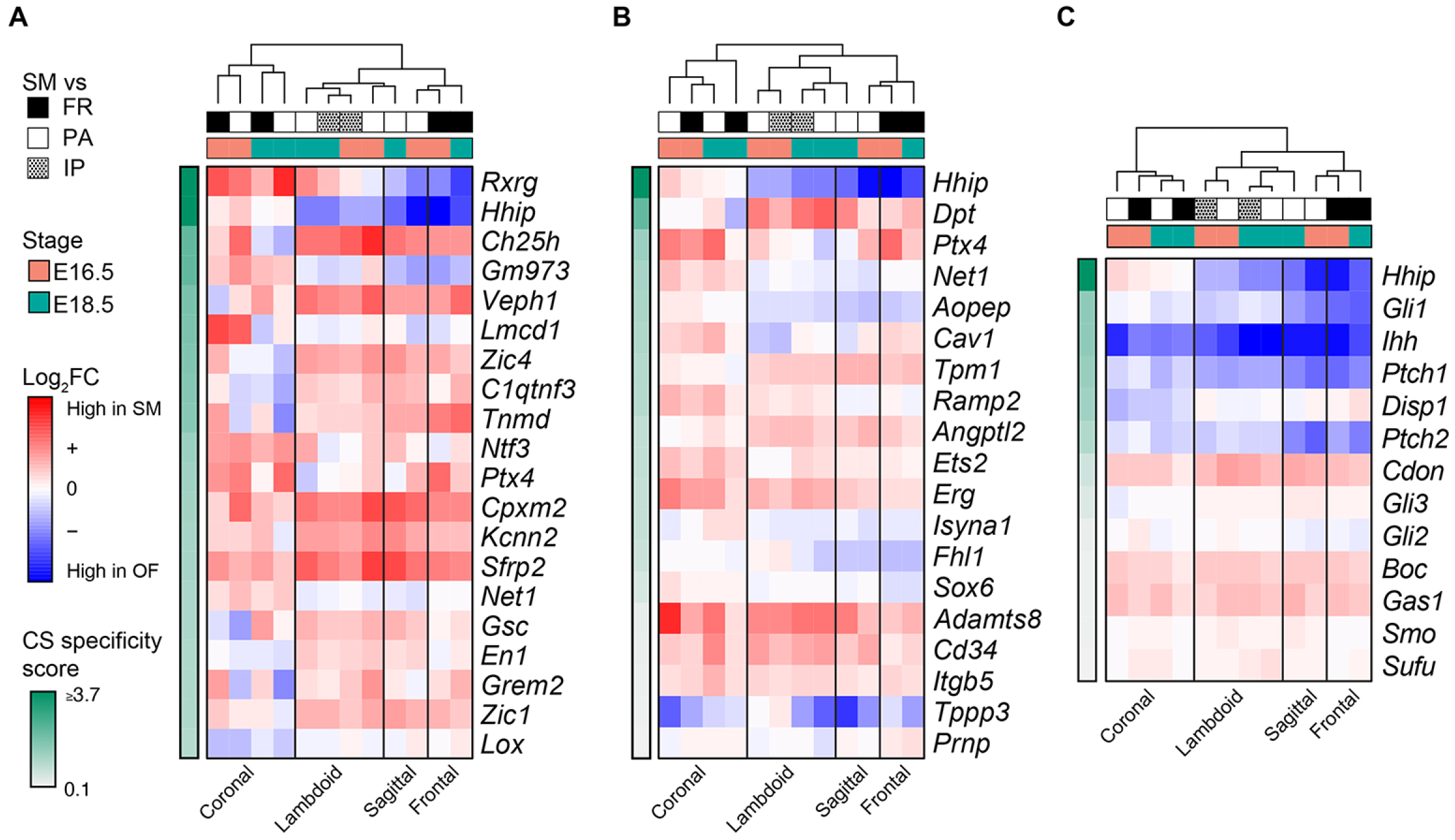
Enrichment of *Hhip* expression in SM is specific to the coronal suture. (*A*) Average bulk gene expression changes of the significant marker genes (rows) of all SM populations across the four calvarial sutures. Expression changes are shown as the log2FC between the SM and osteogenic fronts (OF) at E16.5 and E18.5, ranked by their ‘coronal specificity score’, representing the difference between the average expression across the coronal suture data and the average expression across the remaining sutures (frontal, sagittal, and lambdoid). Frontal OF, FR; parietal OF, PA; interparietal OF, IP. (*B*) Average bulk gene expression changes of the combined top 10 marker genes of SM populations CS6-4 and CS8-2 across the four calvarial sutures. Expression changes are shown as in (*A*). (*C*) Average bulk gene expression changes of key hedgehog pathway genes across the four calvarial sutures. Expression changes are shown as in (*A*).

### *Hhip* is required for normal coronal suture development

We reasoned that *Hhip* expression in the SM is necessary for normal coronal suture development. We examined the coronal suture in *Hhip^-/-^* mice and found a previously unreported coronal suture defect (Fig. 5). At E16.5, staining for alkaline phosphatase (ALPL) activity, expressed in preosteoblasts and osteoblasts, showed that the overlap of frontal and parietal bones seen in the WT is reduced or absent in mutant sutures so that the OFs are more closely apposed (Fig. 5*A*). In the WT suture, RUNX2, a marker of osteoprogenitors and more differentiated osteoblasts, was expressed in the SM, OFs, and more differentiated osteoblasts (Fig. 5*B*) while SP7, a marker of committed preosteoblasts, was expressed in the OFs and more differentiated osteoblasts (Fig. 5*C*). RUNX2 and SP7 expression in mutant sutures localized to the same sutural regions as in WT, despite the altered morphology of the bones (Fig. 5 *B* and *C*). No differences in proliferation of frontal or parietal OFs or SM was seen between WT and mutant sutures (Fig. 5 *D* and *E*). Total cell numbers in the frontal and parietal OFs were similar between WT and mutant sutures, but there was a trend toward lower SM cell numbers in mutant sutures (Fig. 5*F*). Assessment of apoptosis by the TUNEL assay found no apoptotic cells in WT or mutant sutures (Fig. 5*G*).

**Fig. 5.**
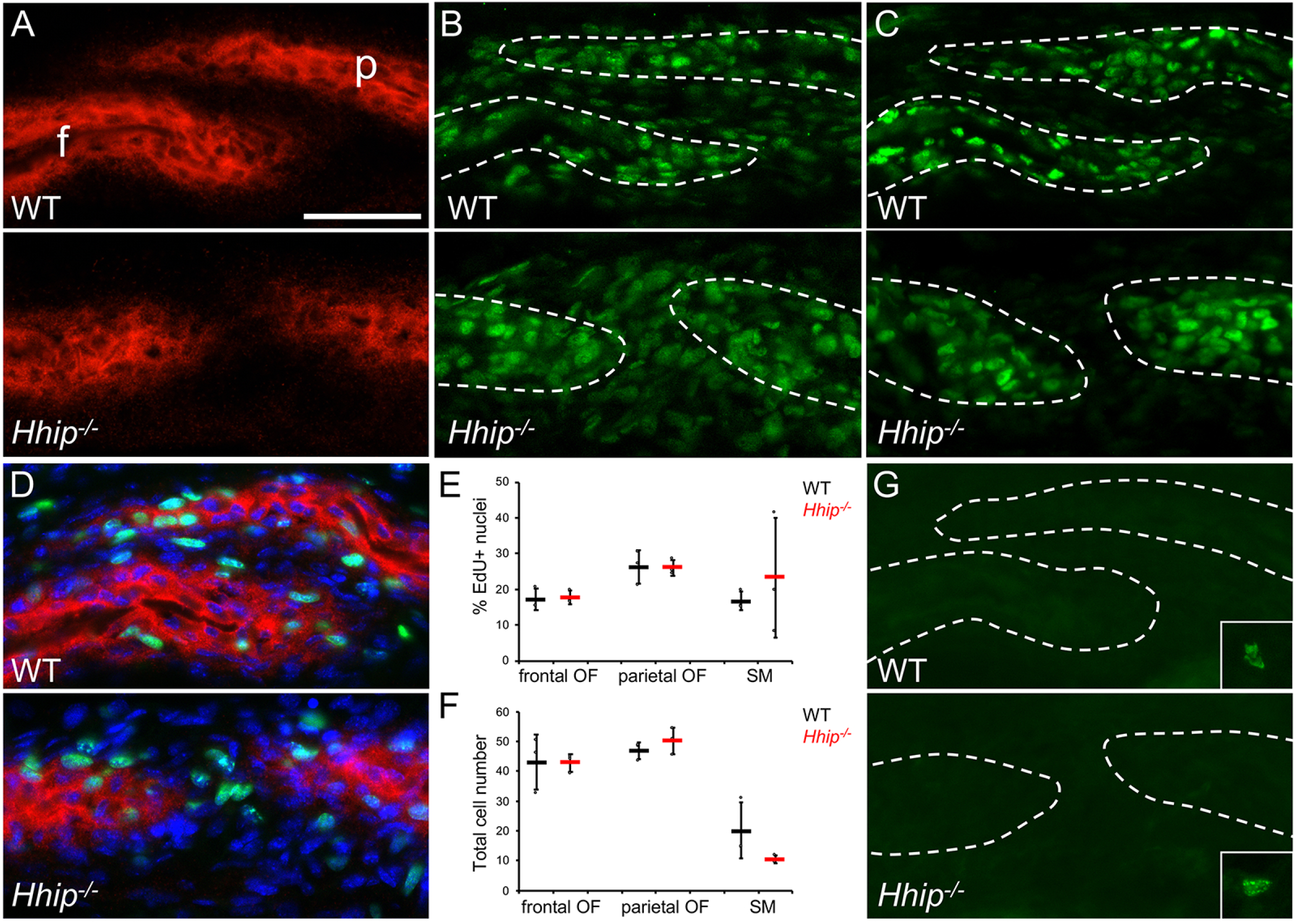
Coronal suture dysgenesis in *Hhip^-/-^* mice at E16.5. (*A*) Staining for alkaline phosphatase activity (ALPL, red) in preosteoblasts and osteoblasts and for nuclei (DAPI, blue) for wild type (WT) and mutant (*Hhip^-/-^*) coronal sutures. f, frontal bone; p, parietal bone. (*B*) Immunohistochemistry for RUNX2 (green). Sections are the same as shown in *A*. (*C*) Immunohistochemistry for SP7 (green). (*D*) Proliferation detected by EdU incorporation (green). Sections are counterstained for ALPL activity (red) and DAPI (blue). (*E*) Quantification of EdU incorporation in osteogenic fronts (OF) and suture mesenchyme (SM). WT and *Hhip^-/-^* are indicated in black and red, respectively. n = 3 WT and 3 *Hhip^-/-^*. (*F*) Quantification of cell numbers in OFs and SM. n = 3 WT and 3 *Hhip^-/-^*. (*G*) TUNEL assay for apoptotic cells. Inset, example of an apoptotic cell in non-sutural tissue on the same slide. White dashed outlines, frontal and parietal bones. Dot plots show mean and standard deviation. All sections are in the transverse plane. Scale bar, 50 μm.

At E18.5 the abnormal *Hhip^-/-^* phenotype was more severe. Coronal sutures still had little or no overlap of frontal and parietal bones along the length of the suture, and in some regions ALPL expression showed that the OFs were so closely apposed that there was little or no intervening SM (Fig. 6*A*). RUNX2 expression in mutant sutures localized to the same sutural regions as in WT (Fig. 6*B*). Where the mutant phenotype was more pronounced, SP7 expression was lower and less distinct between the OFs and presumptive SM than in WT sutures (Fig. 6*C*). A trend toward decreased preosteoblast proliferation in mutant frontal and parietal OFs compared to WT was seen but did not reach significance (Fig. 6 *D* and *E*). Total cell numbers in the mutant frontal and parietal OFs were significantly increased compared to WT, while there was now a significant decrease in SM cell numbers in mutant sutures (Fig. 6*F*). Assessment of apoptosis by the TUNEL assay found no apoptotic cells in WT or mutant sutures (Fig. 6*G*). At P0, ALPL staining showed that cells expressing ALPL activity were continuous between frontal and parietal bones in the most severely affected regions of the coronal suture (*SI Appendix*, Fig. S3). However, neonatal lethality of *Hhip^-/-^* mice due to lung hypoplasia precluded observation of later phenotypes (13). *Hhip* function is clearly required for normal coronal suture development.

**Fig. 6.**
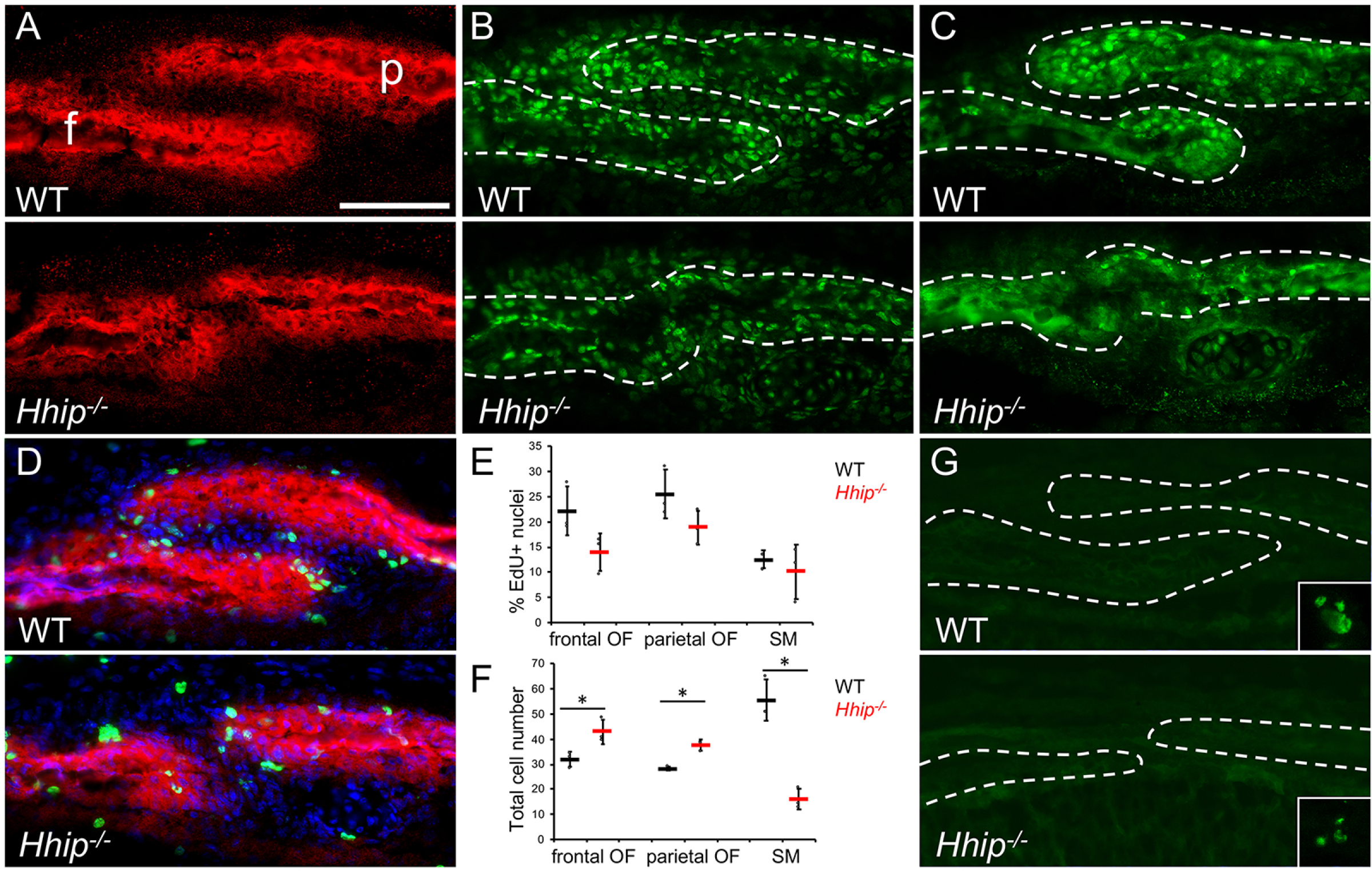
Coronal suture dysgenesis in *Hhip^-/-^* mice at E18.5. (*A*) Staining for alkaline phosphatase activity (ALPL, red) in preosteoblasts and osteoblasts and for nuclei (DAPI, blue) for WT and *Hhip^-/-^* coronal sutures. f, frontal bone; p, parietal bone. (*B*) Immunohistochemistry for RUNX2 (green). Sections are the same as shown in *A*. (*C*) Immunohistochemistry for SP7 (green). (*D*) Proliferation detected by EdU incorporation (green). Sections are counterstained for ALPL activity (red) and DAPI (blue). (*E*) Quantification of EdU incorporation in osteogenic fronts (OF) and suture mesenchyme (SM). WT and *Hhip^-/-^* are indicated in black and red, respectively. n = 3 WT and 3 *Hhip^-/-^*. (*F*) Quantification of cell numbers in OFs and SM. Asterisk indicates a significant difference using the two-tailed Student’s *t* test (*P* = 0.030, 0.002, and 0.002, respectively). n = 3 WT and 3 *Hhip^-/-^*. (*G*) TUNEL assay for apoptotic cells. Inset, example of an apoptotic cell in non-sutural tissue on the same slide. White dashed outlines, frontal and parietal bones. Dot plots show mean and standard deviation. All sections are in the transverse plane. Scale bar, 100 μm.

## Discussion

Using single-cell with bulk RNA-seq we have been able to better define the distinctive composition of the coronal suture at the transcriptional and cell population levels. By scRNA-seq we identified five and eight major cell populations with mesenchymal or osteoblast identities present at E16.5 and E18.5, respectively; these do not include hypodermis or dura mater populations. There were five populations common to both ages: *Hhip*-expressing SM (CS6-4 and CS8-2), ectocranial SM (CS6-6 and CS8-7), proliferating preosteoblasts at both parietal and frontal OFs (CS6-5 and CS8-1), and differentiated *Ibsp*-and *Mmp13*-expressing osteoblasts (CS6-1 and CS8-9; CS6-3 and CS8-5, respectively). The differences between the two ages were the addition at E18.5 of one mesenchyme population and two osteoblast populations. The mesenchyme between the overlapping frontal and parietal bones (CS8-2) was distinguished from the mesenchyme extending beyond this overlap, adjacent to osteogenic fronts (CS8-4). Within osteoblast populations, a distinct periosteal population (CS8-3) and a more differentiated *Spp1*-expressing population were present (CS8-8).

Moreover, populations found in the frontal and parietal bones may differ in their gene signatures because of the division of the coronal suture between the frontal bone derived from neural crest and the parietal bone and SM derived from mesoderm. By comparing the spatial locations of populations with respect to the known boundary of neural crest and mesoderm, we found that the *Hhip*-expressing SM populations (CS6-4, CS8-2, and CS8-4) and ectocranial mesenchyme (CS6-6 and CS8-7) are located in tissue of the mesoderm lineage. The possibility of lineage-specific expression differences also are suggested by previous studies showing that neural crest has increased osteogenic capacity compared to mesoderm. Postnatal frontal bone has an enhanced osteogenic capacity compared to the parietal (24, 25) and embryonic and postnatal frontal bone has increased expression of pro-osteogenic FGF ligands and receptors compared to the parietal (26). Our analysis did identify such a difference in populations between frontal and parietal bones. This was the presence of an *Mmp13*-expressing osteoblast population principally on the endocranial face of the frontal bone (CS6-3 and CS8-5), which was not present on the parietal bone near the suture. This could represent a lineage-specific difference. However, this may be a difference in bone formation processes between frontal and parietal bones in the suture region because *Mmp13*-expressing osteoblasts, known to be involved in bone remodeling (27, 28), were present in more mature regions of both bones away from the suture. Expression of *Mmp13* also has been reported in the human embryonic calvaria in more mature endocranial and trabecular bone (29). Indeed, we did not identify lineage-specific distinctions for other populations common to neural crest-and mesoderm-derived tissue. We found only one proliferating osteoblast population that localized to both OFs at each age (CS6-5 and CS8-1). Similarly, the *Ibsp*-, *Spp1*-, and *Postn*/*Col6a3*-expressing osteoblast populations (CS6-1, CS8-9, CS8-8, and CS8-3) are present in both frontal and parietal bones.

Many genes have been identified that cause coronal and other calvarial suture fusion when mutated in humans. In particular, the ectocranial mesenchyme population present at both ages and the periosteal and *Ibsp*-expressing osteoblast populations population at E18.5 were enriched for such genes. Significant enrichment also was found in the *Mmp13*-expressing population, but as this appears to be a specialized, differentiated osteoblast population, some caution must be used in interpreting this finding. At one or both ages, this population includes *Alpl*, *Fgfr2*, and *Fgfr3* as marker genes. While these may be more highly expressed in the *Mmp13*-expressing population in comparison to other populations, these genes are still expressed in the OFs and osteoblasts. Mutations in these genes therefore may exert their influence through other cell populations. In addition, we have assessed the coronal suture populations at E16.5 and E18.5, but mutated genes may act on suture development if expressed at earlier developmental stages.

We noted differences between the coronal and other calvarial sutures. Coronal suture cell populations differed from the frontal suture, which we previously characterized by single-cell and bulk RNA-seq (12). The major differences were between the SM populations of each suture. *Hhip* expression was a feature of coronal suture SM at E16.5 and E18.5, while its expression was enriched in OFs in the frontal suture at these ages. A mesenchymal population in the frontal suture, enriched for expression of *Acta2* and other genes involved in cell contractility or tendon and ligament development, was not found in the coronal suture. This suggests differences in the mechanical environment of frontal and coronal sutures in which frontal bones are apposed end-to-end in the frontal suture, while frontal and parietal bones overlap in the coronal suture. This differing morphology may be influenced by or shape responses to potential differences in tension forces at each suture during expansion of the skull and brain. Finally, the *Mmp13*-expressing osteoblast population was not identified in the frontal suture.

Enriched expression of *Hhip*, an inhibitor of HH signaling, in coronal SM suggests a difference in the regulation of suturogenesis by HH signaling between coronal and other calvarial sutures. A comparison of single-cell SM gene expression to bulk RNA-seq data from all four calvarial sutures confirmed that *Hhip* enrichment in SM was unique to the coronal suture. In contrast, SM-specific enrichment of other SM marker genes such as *Cpxm2* and *Sfrp2* was common to all four sutures. A similar comparison of key HH pathway genes also identified *Hhip* as the only pathway member with expression specifically enriched in the SM of the coronal compared to other calvarial sutures.

HH signaling plays an important role in embryonic suturogenesis, where it is believed to have pro-osteogenic functions in the OFs and SM (30). *Ihh* is expressed in OFs and is believed to be the only functional HH ligand during calvarial development (31). In *Ihh*^-/-^ calvaria, intramembranous ossification is decreased resulting in wider sutures (31–34). In both humans and mice, craniosynostosis affecting various sutures including the coronal results from the duplication of regulatory elements driving *Ihh* expression (35–37). Mutations in other HH pathway members such as *Ptch1*, *Smo*, *Gli3*, and *Rab23* also cause craniosynostosis in humans and/or mice (38–41). HH signaling is also important for the function of a suture stem cell population that develops postnatally in all craniofacial sutures and is responsible for maintenance and repair of calvarial bone (42).

Our finding of *Hhip* expression in SM populations (CS6-4 and CS8-2) suggests that inhibition of functional HH signaling is required in SM at least in later embryonic stages for normal suturogenesis. Indeed, we found that loss of *Hhip* results in altered coronal suture morphology by E16.5 and depletion of SM at E18.5 and P0, so that OFs were closely apposed or even bridged by osteogenic cells expressing ALPL. We did not find differences in proliferation or apoptosis between WT and mutant sutures to explain this defect, but by E18.5 expression of SP7 was lower in intensity compared to WT in regions of the suture where OFs were most closely apposed. Taken together, this suggests that the regulation of osteoprogenitor recruitment from SM is altered in neonatal lethal, *Hhip^-/-^* mutants leading to loss of SM and potential suture fusion.

During embryonic suture development there is essentially no mixing between neural crest and mesoderm lineages within the coronal suture (9, 10). Osteoprogenitors forming the neural crest-derived frontal bone must be restricted to the region of neural crest-derived mesenchyme at the OF. In contrast, the mesoderm-derived parietal bone and SM are contiguous, and so the SM potentially provides a large pool of osteoprogenitors along the parietal bone surface. However, the presence of similar proliferating osteoprogenitor populations at the frontal and parietal OFs suggests a similar mechanism for spatially limiting recruitment of osteoprogenitors to the OFs that may be independent of cell lineage. *In vivo* labeling studies of the frontal and sagittal sutures also suggest that bone growth is sustained by the proliferation of osteoprogenitors within the OFs with minor recruitment of adjacent SM cells (10, 43). As RUNX2 is expressed in cells throughout the coronal SM, these are all potential osteoprogenitors, and so must be excluded from the mechanism of osteoprogenitor recruitment. This mechanism could in part consist of IHH expressed by osteoblasts at the OFs (CS6-1 and CS8-9) acting on an adjacent pool of osteoprogenitors (CS6-5 and CS8-1). Osteoprogenitor recruitment from the SM between the frontal and parietal bones would be inhibited by the presence of HHIP to maintain a non-ossifying suture mesenchyme. HH signaling is not likely to be the sole mechanism regulating this process, but would interact with pathways such as FGF and BMP that also regulate the balance between osteoprogenitor recruitment and preservation of suture mesenchyme (32, 44).

In conclusion, we have transcriptionally defined cell populations of the murine coronal suture and their constituent marker genes during embryonic development using scRNA-seq. We found that enriched *Hhip* expression was a feature of a specific SM population distinguishing the coronal suture from other calvarial sutures. This discovery led us to identify a novel coronal suture phenotype for *Hhip^-/-^* mice, in which potential osteoprogenitors are depleted from the SM. We propose a revised view of the role of HH during suturogenesis in which functional HH signaling is excluded from the SM by HHIP. Our approach greatly expands opportunities for hypothesis-driven research in coronal and other suture development.

## Materials And Methods

### Mice

Mouse procedures were in compliance with animal welfare guidelines mandated by the Institutional Animal Care and Use Committee (IACUC) of the Icahn School of Medicine at Mount Sinai. Timed matings of C57BL/6J mice (The Jackson Laboratory, 000664) or *Hhip^tm1Amc^*/J mice (The Jackson Laboratory, 006241; homozygotes referred to as *Hhip^-/-^*) were performed to obtain embryos at required ages. Genotyping was performed by polymerase chain reaction (PCR) of tail DNA. Sex genotypes were identified by PCR as described previously (45, 46). *Hhip* genotypes were identified by PCR as described by The Jackson Laboratory.

### scRNA-seq library preparation

Coronal sutures, encompassing the SM, OFs, and frontal and parietal bones in the region of overlap, were dissected from C57BL/6J mice (Fig. 1*A*). Ectocranial and endocranial membranes were removed to the extent possible, and frontal and parietal bones were separated with forceps to expose the SM. Sutures of both sexes were combined for processing. Two libraries each at E16.5 and E18.5 were created. The E16.5 libraries each consisted of pooled sutures from 13 embryos obtained on the same day; the E18.5 libraries each consisted of pooled sutures from 8 and 13 embryos obtained on separate days. Pooled E16.5 sutures were digested at 37°C in αMEM (Gibco, 32571-036) with 0.2% collagenase type II (Worthington, LS004176), 0.2% dispase II (Sigma-Aldrich, 4942078001), and 1U/μl DNase (Qiagen, 79254). Sutures were serially digested with agitation five times for 10 minutes each in a shaking incubator. Successive fractions were pooled on ice with the addition of FBS to 2%. Cell suspensions were strained through a 40 μm filter (Falcon, 352340), pelleted at 400g for 7 minutes, washed in PBS/1% BSA, and resuspended in PBS/1% BSA. Pooled E18.5 sutures were similarly processed and red blood cell lysis (Miltenyi Biotec, 130-094-183) was performed after pelleting filtered cells. scRNA-seq 3’ expression libraries were prepared on a Chromium instrument (10X Genomics, model GCG-SR-1) using the Chromium Single Cell Gene Expression kit (version 3 for E16.5 libraries; version 2 for E18.5 libraries) by the Technology Development Facility at the Icahn School of Medicine at Mount Sinai. Sequencing was conducted by the Genetic Resources Core Facility, Johns Hopkins Institute of Genetic Medicine. Libraries were first run on an Illumina MiSeq at 26×8×98 and analyzed to confirm the number of captured cells and assess capture efficiency prior to sequencing. E16.5 libraries were sequenced in a single pool on an Illumina NovaSeq S1. E18.5 libraries were sequenced separately on an Illumina HiSeq 2500. Sequencing used standard Illumina primers, where read 1 was the UMI, read 2 was the library index, and read 3 was the transcript. E16.5 libraries consisted of a final number of 941 and 1,185 cells. E18.5 libraries consisted of a final number of 2,625 and 1,624 cells.

### scRNA-seq analysis

Preprocessing of 10X Genomics scRNA-seq data, clustering and cell type identification was done as previously described with a few modifications (12). Cells with >800 genes detected, and genes detected in >5 cells were included. The cells from both replicates and both embryonic days were normalized using regularized binomial negative regression (47) and integrated based on common transcripts or ‘anchors’ (48). The cleaned and integrated data contained 19,903 genes and 5,142 cells across all samples, and was then used for initial clustering. Thirteen PCs were used to perform unsupervised shared nearest neighbor graph-based clustering (k = 100) as implemented in the package Seurat (48, 49). Differential gene expression analysis among cell clusters and stages was tested by logistic regression using replicate as a latent variant, an FDR ≤ 0.05 and lnFC expression threshold ≤ 0.25. Mitochondrial and erythroid-specific gene expression were treated as covariates as previously described (12).

### Identification of subpopulations within the coronal suture

To analyze the relationship between the suture mesenchyme, osteoblasts and adjacent populations in the initial UMAP plot, we performed a second unsupervised clustering (k parameter = 50, resolution = 0.8) and differential gene expression analysis for these cell types separately for each embryonic day, using the 13 and 15 top PCs for E16.5 and E18.5 respectively. In addition, we used a combined Fisher p value of 0.01 and a minimum lnFC threshold of 0.1 to determine significant markers (50).

### Gene ontology enrichment analyses

Gene ontology (GO) biological process (BP), molecular function (MF), and/or cellular component (CC) enrichment analyses for each single cell cluster were performed using the gProfileR R v0.6.4 package (51) as previously described (12).

### Gene enrichment analysis

Lists of 96 human and 31 mouse genes associated with craniosynostosis and subsets of genes associated with coronal synostosis (5, 7, 22, 23) (*SI Appendix*, Table S2) were tested for significant (p ≤ 0.05) gene set enrichment against the single cell CS populations, determined by scRNA-seq analysis using Fisher’s exact test and using Bonferroni correction for multiple comparisons. This method was also used to match the single cell CS populations identified at E16.5 with those identified at E18.5.

### Comparative analysis of calvarial suture RNA-Seq data

To compare gene expression profiles of calvarial sutures we leveraged RNA-Seq data that we generated for SM and OF regions of 11 craniofacial sutures as part of the “Transcriptome Atlases of the Craniofacial Sutures” FaceBase2 project (52, 53). Briefly, count-per-million (CPM) values for 629 SM and OF RNA-seq libraries for coronal, frontal, sagittal, lambdoid, intermaxillary, internasal, interpremaxillary, interpalatine, maxillary-palatine, premaxillary-maxillary, and squamoparietal sutures at E16.5 and E18.5 (up to 5 replicates each) were filtered to retain genes with ≥1 CPM in more than 10% of samples, and then normalized using the weighted trimmed mean of M-values (TMM) method (54). Finally, we applied the Voom transformation (54) to the data, which included transforming the matrix to log2 CPM data, estimating the mean-variance relationship and computing appropriate observation-level weights using this relationship. Differential gene expression analysis was then performed between the OF and SM regions of the calvarial sutures as described previously (12).

### Alkaline phosphatase (ALPL) staining

ALPL staining was performed as described previously (12). Nuclei were stained with DAPI.

### Immunohistochemistry and cytochemistry

Hematoxylin and eosin stained sections were prepared from paraffin-embedded calvaria by standard procedures. Antibody staining for RUNX2 (1:200; rabbit anti-RUNX2, Sigma-Aldrich, HPA022040) and SP7 (1:500; rabbit anti-SP7/Osterix, Abcam, ab22552) was performed after ALPL staining using standard procedures. Primary antibodies were detected with donkey anti-rabbit IgG Alexa Fluor 488 (1:400; Thermo Fisher Scientific, A-21206). For EdU quantification, pregnant mice were injected with EdU (250 μg/10 kg body weight) two hours before sacrifice. EdU staining was performed with the Click-iT Plus EdU Alexa Fluor 488 Imaging Kit (Thermo Fisher Scientific, C10637) as described by the manufacturer. Immunohistochemistry and EdU staining were performed on 10 μm sections from either fresh-frozen or 4% paraformaldehyde-fixed cryoembedded heads prepared as previously described (55). Sections were stained with DAPI. EdU-positive nuclei and total nuclei were counted within the OFs and SM. In OFs nuclei were counted within the ALPL-positive domain between the SM and the start of the osteoid, or within a 50 μm distance if the osteoid was not apparent. SM was defined as ALPL-negative cells between the ends of the ALPL-positive OFs. Cell counts were performed in Adobe Photoshop. TUNEL staining was performed using the In Situ Cell Death Detection Kit, Fluorescein (Roche, 11684795910) as described by the manufacturer. Images of histological sections were taken using a Nikon Eclipse E600 microscope equipped with a Nikon DS-Ri2 digital camera and NIS Elements (F4.30.01) software.

### Single molecule fluorescent RNA *in situ* hybridization (smFISH)

smFISH was performed using the RNAscope Fluorescent Multiplex Reagent Kit (Advanced Cell Diagnostics, 320850) with modifications as described previously (12). Probes (Advanced Cell Diagnostics) were for *Col6a3* (552541-C2), *Col8a1* (518071), *Clec3b* (539561-C2), *Igfbp3* (405941*), Cxcl12* (422711-C3), *Erg* (546491), *Fxyd5* (527721-C2), *Hhip* (448441-C3), *Ibsp* (415501), *Mest* (405961), *Mmp13* (427601-C3), *Postn* (418581-C2), *Sox6* (472061-C2), *Spp1* (435191-C3), and *Top2a* (491221). Images were acquired on an AxioImager Z2M equipped with a 20x/0.8NA Zeiss Plan-Apochromat objective, a monochrome Axiocam 503 camera (Zeiss, 1936 × 1460 pixels, 4.54 µm X 4.54 µm per pixel, sensitivity ∼400 nm–1000 nm) and Zen 2 Blue Edition software (version 2.0). Z-stack images were acquired at optimal sampling rate meeting Nyquist frequency requirements, as calculated by the software.

### Statistical Analysis

For quantification of EdU incorporation and cell number, statistical analysis and dot plotting were performed with Microsoft Excel for Mac, version 16.16.27. Dot plots show the mean and standard deviation. Statistical significance was determined using the two-tailed Student’s *t* test. *P* values below 0.05 were considered significant.

### Data Availability

The data for the single-cell RNA-seq (accession FB00000970) and bulk RNA-seq libraries (accession no. FB00000903, FB00000902, FB00000805, FB00001076, and FB00000998) reported in this study are part of “Transcriptome Atlases of the Craniofacial Sutures” FaceBase2 project, and available in the FaceBase data repository (facebase.org).

## ACKNOWLEDGMENTS

This work was funded by NIH grants U01DE024448 (FaceBase Spoke grant to G.H., H.v.B., and E.W.J.), R03DE026814 (to B. Z. and E.W.J.), and P01HD078233 (to E.W.J.). This work was supported in part through the computational resources and staff expertise provided by Scientific Computing at the Icahn School of Medicine at Mount Sinai.

## Notes

The authors declare no conflict of interest.

### Competing Interest Statement

The authors have declared no competing interest.

